# Mitochondria promote neuropeptide secretion in *Caenorhabditis elegans* by preventing activation of hypoxia inducible factor

**DOI:** 10.1101/298034

**Authors:** Tongtong Zhao, Joshua M Kaplan

## Abstract

Neurons are highly dependent on mitochondrial function, and mitochondrial damage has been implicated in many neurological and neurodegenerative diseases. Relatively little is known about how mitochondria regulate neuronal function. Here we show that axonal mitochondria are necessary for neuropeptide secretion in *Caenorhabditis elegans*, and that oxidative phosphorylation, but not mitochondrial calcium uptake, is required for secretion. Oxidative phosphorylation produces cellular ATP, reactive oxygen species, and consumes oxygen. Disrupting any of these functions could inhibit neuropeptide secretion. We show that blocking mitochondria transport into axons inhibits neuropeptide secretion through activation of the hypoxia inducible factor HIF-1. Our results suggest that axonal mitochondria modulate neuropeptide secretion by regulating transcriptional responses induced by metabolic stress.

## INTRODUCTION

Mutations that perturb mitochondrial function are linked to several neurodevelopmental and neurodegenerative disorders (Johri and Beal 2012; Pease and Segal 2014; Pickrell and Youle 2015; Kerr *et al*. 2017; Krench and Littleton 2017). For these reasons, there is significant interest in defining the cell biological mechanisms through which mitochondria regulate neuronal function and development. Mutations (or drug treatments) that impair mitochondrial function cause widespread cellular and developmental defects. Given these pleiotropic effects, identifying the precise molecular pathways through which mitochondria regulate neural development and behavior has proved challenging. One strategy to circumvent this problem has been to analyze mutants in which mitochondria transport or localization in axons is blocked (Guo *et al*. 2005; Verstreken *et al*. 2005; Kang *et al*. 2008; Ma *et al*. 2009; Russo *et al*. 2009). In these transport mutants, mitochondrial defects are restricted to axons, whereas other cellular functions of mitochondria remain intact.

Analysis of mitochondria transport mutants suggests mitochondria can act through multiple pathways to influence neuronal function and development. Several aspects of neuronal function are especially dependent upon cellular energy stores (e.g. maintenance of membrane potential, intracellular transport to axons and dendrites, and membrane fusion and endocytosis reactions). In addition to producing ATP, mitochondria also act as an intracellular calcium store; consequently, axonal mitochondria alter the duration of presynaptic calcium transients thereby regulating neurotransmitter release at synapses (Werth and Thayer 1994; Tang and Zucker 1997; Billups and Forsythe 2002; Medler and Gleason 2002). Finally, mitochondria promote axon regeneration following injury (Rawson *et al*. 2014; Han *et al*. 2016). Given the widespread effects of mitochondria on cellular physiology, it is likely that additional mechanisms will be discovered that link mitochondria to neuronal function and development.

To further investigate how they impact neuronal function, we asked if axonal mitochondria regulate secretion of neuropeptides. Neuropeptides are a chemically diverse set of neuromodulators that are recognized by G protein coupled receptors (GPCRs). GPCRs typically bind their cognate ligands with extremely high affinity and stimulate production of second messengers through G-protein activation of enzymatic targets. Consequently, neuropeptides (and other metabotropic agonists) often produce relatively long-lasting signals over very large spatial domains. Thus, we hypothesized that axonal mitochondria could alter global behavioral states by regulating neuropeptide secretion. Here we show that axonal mitochondria promote neuropeptide secretion and that they do so by regulating a cell-autonomous stress response to hypoxia.

## MATERIALS AND METHODS

### C. elegans strains and drug assays

Strains were maintained at 20°C as described (Brenner 1974). The wild-type reference strain was N2 Bristol. Full descriptions of all alleles can be found at http://www.wormbase.org. Strains used in this study are as follows:

KP8711 *ric-7(nu447) V*

KP8914 *nuEx1898[punc-129::SNB-1-GFP, punc-129::TOMM-20-mCherry]*

KP8730 *ric-7(nu447) V; nuEx1853[psnb-1::UNC-116-GFP-TOMM-7]*

KP8717 *nuIs183 III; nuEx1851[punc-129::UNC-116-RFP-TOMM-7]*

KP8723 *ric-7(nu447) V; nuIs183 III; nuEx1851[punc-129::UNC-116-RFP-TOMM-7]*

KP9082 *nuSi187 IV; nuEx1851[punc-129::UNC-116-RFP-TOMM-7]*

KP9083 *ric-7(nu447) V; nuSi187 IV; nuEx1851[punc-129::UNC-116-RFP-TOMM-7]*

KP8755 *nuIs183 III; mcu-1(tm6026) IV*

KP8756 *nuIs183III; mcu-1(ju1154) IV*

KP8749 *nuIs183 III, mev-1(kn1) III*

KP8748 *nuIs183 III; isp-1(qm150) IV*

KP8759 *nuo-6(qm200) I; nuIs183 III*

KP8777 *nuo-6(qm200) I; nuIs183 III; nuEx1863[punc-129::NUO-6]*

KP8803 *sod-1(tm776) II; nuIs183 III*

KP8804 *sod-2(ok1030) I; nuIs183 III*

KP8874 *sod-2(ok1030) I; nuIs183 III; nuEx1887[punc-129::SOD-2]*

KP8839 *sod-3(tm760) X; nuIs183 III*

KP8831 *nuIs183 III, sod-4(gk101) III*

KP8832 *sod-5(tm1146) II; nuIs183 III*

KP9044 *nuSi187 IV*

KP9073 *nuSi187 IV; ric-7(nu447) V*

KP9099 *sod-2(ok1030) I; nuSi187 IV*

KP9100 *sod-2(ok1030) I; nuSi187 IV; ric-7(nu447) V*

KP8778 *vhl-1(ok161) X; nuIs183 III; nuEx1864[punc-129::VHL-1]*

KP8780 *vhl-1(ok161) X; nuIs183 III; nuEx1866[pvhl-1::VHL-1]*

KP9109 *vhl-1(ok161) X; nuSi187 IV*

KP9110 *vhl-1(ok161) X; nuSi187 IV; ric-7(nu447)*

KP9111 *vhl-1(ok161) X; nuSi187 IV; hif-1(ia4) V*

KP9067 *nuIs183 III, aldo-1(tm5782) III*

KP9069 *pfkb-1.1(ok2733) I; nuIs183 III*

KP9071 *gpd-3(ok2870) X; nuIs183 III*

KP9097 *nuSi187 IV*; *hif-1(ia4) V*

KP9098 *nuSi187 IV*; *ric-7(nu447) V, hif-1(ia4) V*

KP9095 *daf-16(mgDf47) I; nuSi187 IV*

KP9096 *daf-16(mgDf47) I; nuSi187 IV; ric-7(nu447) V*

KP9106 *nuSi187 IV, unc-31(e928) IV*

KP9112 *vhl-1(ok161) X; nuSi187 IV, unc-31(e928) IV*

### Constructs and transgenes

Transgenic strains were isolated by microinjection of various plasmids using either P*myo-2*::NLS-GFP or P*myo-2*::NLS-mCherry as co-injection markers. The mito-truck (mtruck) construct was adapted from the original version (Rawson *et al*. 2014) to contain the following elements (listed 5’ to 3’): the *snb-1* promoter (for locomotion and aldicarb assays) or the *unc-129* promoter (for imaging), the cDNA for *unc-116*, linker sequence, GFP (for locomotion and aldicarb) or TagRFP (for imaging) with syntrons, att recombination site + second linker, the genomic sequence of *tomm-7*, and the *let-858* 3’UTR.

The TOMM-20-mCherry construct contains the *unc-129* promoter, the sequence encoding the first 53 amino acids of TOM-20 carrying the mitochondrial localization signal, att recombination site, mCherry with syntrons, and the *unc-54* 3’UTR. The P*unc-129::*SNP-1- GFP construct was previously described (Sieburth *et al*. 2005).

For the P*unc-129::*NLP-21-mNeonGreen construct the *unc-129* promoter, the *nlp-21*- minigene (coding region plus two syntrons), mNeonGreen with syntrons, and the *let-858* 3’UTR were inserted into the miniMOS vector pCFJ910. Single copy insertion *(nuSi187*) of this transgene was obtained as described (Frokjaer-JENSEN *et al*. 2014). The P*unc-129::*NLP-21-YFP (*nuIs183*) transgenic line was previously described (Sieburth *et al*. 2007).

The cell-specific rescue constructs contain the *unc-129* promoter, the open reading frame including introns (*nuo-6*, *vhl-1*) or the cDNA (*sod-2*) of the rescuing gene, and the *unc-54* 3’UTR. The *vhl-1* genomic rescue construct contains the genomic *vhl-1* locus spanning the region 1.5 kb upstream of the start codon to 0.8 kb downstream of the stop codon.

### Drug, hypoxia, and RNAi assays

Acute aldicarb assays were performed blind in five replicates on young adult worms as described (Nurrish *et al*. 1999). Locomotion was measured by transferring adults to unseeded plates and recording their movement at 2 Hz for 30 s on a Zeiss Discovery Steromicroscope using Axiovision software. The centroid velocity of each animal was analyzed at each frame using object-tracking software in Axiovision. The speed of each animal was calculated by averaging the velocity value at each frame.

For treatment of worms with varying oxygen levels, worms were picked onto a fresh plate at the L4 stage and the plate was put into a Hypoxia Incubator Chamber (STEMCELL Technologies). Using a MINIOS 1 oxygen analyser (MEDIQUIP), the chamber was then filled with oxygen (for hyperoxia) or nitrogen until the oxygen test concentration was reached. Worms were left to grow at control and test oxygen levels for 24 hours and coelomocytes were imaged subsequently.

RNAi was performed in the neuronal RNAi hypersensitive mutant background (*nre-1 lin-15b*) (Schmitz *et al*. 2007). One-day old hermaphrodites were dropped into bleach on RNAi plates. When the progeny reached L4 stage, worms were transferred onto a fresh RNAi plate and scored for coelomocyte fluorescence 24 hours later.

### Fluorescence microscopy and quantitative analysis

Worms were immobilized on 10% agarose pads with 0.1 um diameter polystyrene 501 microspheres (Polysciences 00876-15, 2.5% w/v suspension). For axonal fluorescence measurements, the dorsal nerve cord (midway between the vulva and the tail) was captured using a 60x objective (NA 1.45) on an Olympus FV-1000 confocal microscope at 5× digital zoom. Maximum intensity projections of z-series stacks were made using Metamorph 7.1 software (Molecular Devices, Sunnyvale, CA, United States). Line scans of dorsal cord fluorescence were analyzed in Igor Pro (WaveMetrics, Lake Oswego, OR, United States) using custom-written software (Burbea *et al*. 2002; Dittman and Kaplan 2006).

For coelomocyte fluorescence measurements, worms were imaged 24 hours after the L4 stage. Image stacks of the second most posterior coelomocyte were captured using a Zeiss Axioskop I, Olympus PlanAPO 100× 1.4 NA objective, and a CoolSnap HQ CCD camera (Roper Scientific, Tuscon, AZ, United States). Maximum intensity projections were obtained using Metamorph 7.1 software (Molecular Devices). For quantification, the five brightest vesicles were analyzed for each coelomocyte and the mean fluorescence for each vesicle was logged. For each worm, coelomocyte fluorescence was calculated as the mean of the vesicle values in that animal subtracted by the background fluorescence (measured at an empty region on the slide). All fluorescence values are normalized to wild type controls to facilitate comparison.

## RESULTS

### Axonal mitochondria promote neuropeptide secretion

In a prior study, we showed that mutations inactivating the *C. elegans ric-7* gene significantly reduced neuropeptide secretion (which is mediated by exocytosis of dense core vesicles, DCVs) but had little effect on acetylcholine release (which is mediated by exocytosis of synaptic vesicles, SVs) (Hao *et al*. 2012). These results imply that RIC-7 plays a specific role in promoting exocytosis of DCVs but not SVs. Decreased neuropeptide secretion in *ric-7* mutants was accompanied by decreased locomotion rate, and increased resistance to the paralytic effects of aldicarb (an acetylcholinesterase inhibitor) (Hao *et al*. 2012). Homologs of *ric-7* are found in other nematodes but not in other metazoans. The *ric-7* gene encodes an unfamiliar protein that lacks any previously described structural domain. Thus, the sequence of the RIC-7 protein failed to provide any clues as to the cell biological mechanism underlying RIC-7’s function in neuropeptide secretion.

A subsequent study showed that RIC-7 is required for kinesin-1 mediated transport of mitochondria into axons (Rawson *et al*. 2014). These results suggest that the behavioral and neuropeptide secretion defects observed in *ric-7* mutants could be caused by the absence of axonal mitochondria. To test this idea, we expressed a chimeric kinesin construct (mtruck) in which UNC-116/KIF5 is fused to the outer mitochondrial membrane protein TOM-7. As previously reported (Rawson *et al*. 2014), the mtruck transgene rescued the *ric-7* mitochondrial transport defect (Fig. 1A, B). Neuronal expression of mtruck significantly improved locomotion and decreased the aldicarb resistance of *ric-7* mutants (Fig. 1C, D), suggesting that *ric-7* behavioral defects are caused by the lack of axonal mitochondria.

**Figure 1:**
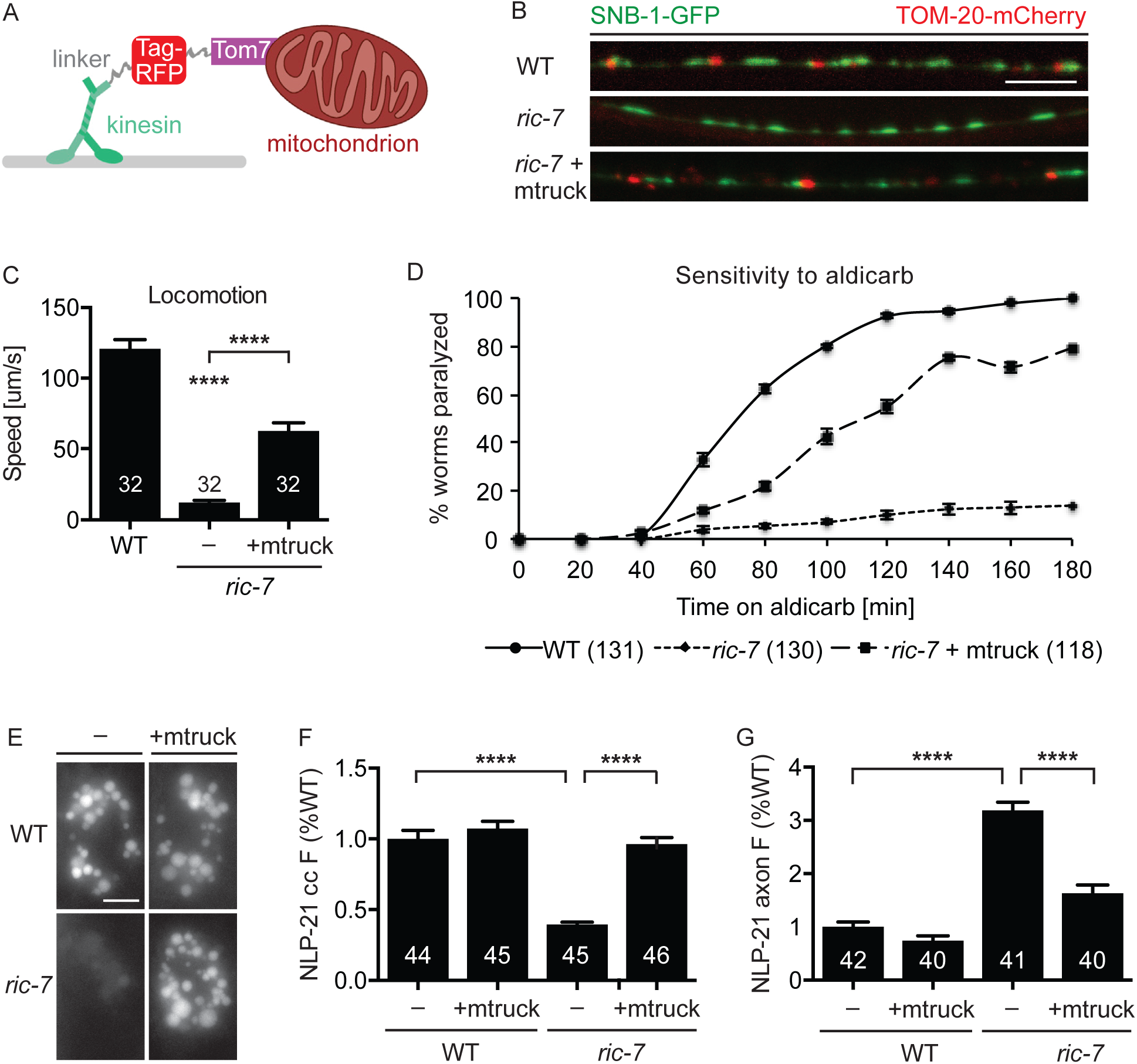
Axonal mitochondria are required for locomotion and neuropeptide release. (**A**) Schematic of the mito-truck (mtruck) construct in which UNC-116/KIF5 was fused to TagRFP and the outer mitochondrial protein TOM-7. This illustration was modified from that shown in the original paper describing mtruck (Rawson *et al*. 2014). (**B**) Localization of the synaptic marker SNB-1 and the mitochondrial marker TOM-20 in dorsal cord axons of cholinergic motor neurons in wildtype animals (top), *ric-7* mutants (middle), and *ric-7* mutants expressing mtruck (bottom). (**C, D**) Locomotion rate (**C**) and aldicarb-induced paralysis (**D**) of wildtype, *ric-7* mutant, and *ric-7* mutant worms that express mtruck in neurons. (**E, F**) Representative images (**E**) and quantification (**F**) of NLP-21 coelomocyte fluorescence in wildtype and *ric-7* mutant worms with and without the DA/DB-specific expression of mtruck. (**G**) Dorsal cord axonal NLP-21 fluorescence in cholinergic motor neurons of wildtype and *ric-7* mutant worms with and without the DA/DB-specific expression of mtruck. Number of animals analyzed is indicated for each genotype. Scale bars are 5 um. Error bars indicate SEM. Values that differ significantly are indicated (****, p < 0.0001, ANOVA).

To assay neuropeptide secretion, we expressed the proneuropeptide NLP-21 tagged with YFP in the cholinergic DA/DB motor neurons (using the *unc-129* promoter). We quantified NLP-21::YFP secretion by analyzing YFP fluorescence in the endolysosomal compartment of coelomocytes, which are scavenger cells that endocytose proteins secreted into the body cavity (Fares and Greenwald 2001). In parallel, we assayed the abundance of NLP-21 containing vesicles in DA/DB axons by quantifying the fluorescence intensity of YFP puncta in dorsal cord axons (Sieburth *et al*. 2007). As previously reported, NLP-21 coelomocyte fluorescence was significantly decreased in *ric-7* mutants while axonal puncta fluorescence was significantly increased (Fig. 1E-G) (Hao *et al*. 2012), both indicating decreased NLP-21 secretion in *ric-7* mutants. Both the coelomocyte and axonal NLP-21 fluorescence defects exhibited by *ric-7* mutants were rescued by expressing mtruck in the DA/DB neurons (Fig. 1E-G). Thus, the *ric-7* behavioral and neuropeptide secretion defects are both caused by the absence of axonal mitochondria.

### Neuropeptide secretion requires oxidative phosphorylation but not mitochondrial calcium uptake

Loss of axonal mitochondria is predicted to result in multiple defects. For example, mitochondria regulate cytoplasmic calcium levels by taking up calcium through the Mitochondrial Calcium Uniporter (MCU) (Kirichok *et al*. 2004; Baughman *et al*. 2011; De STEFANI *et al*. 2011), which could account for the *ric-7* neuropeptide secretion defect. To test this idea, we analyzed two independent *mcu-1* null mutants, finding no significant changes in NLP-21 coelomocyte and axonal puncta fluorescence (Fig. 2A, B). These results suggest that changes in mitochondrial calcium uptake are unlikely to account for the *ric-7* mutant neuropeptide secretion defects.

**Figure 2:**
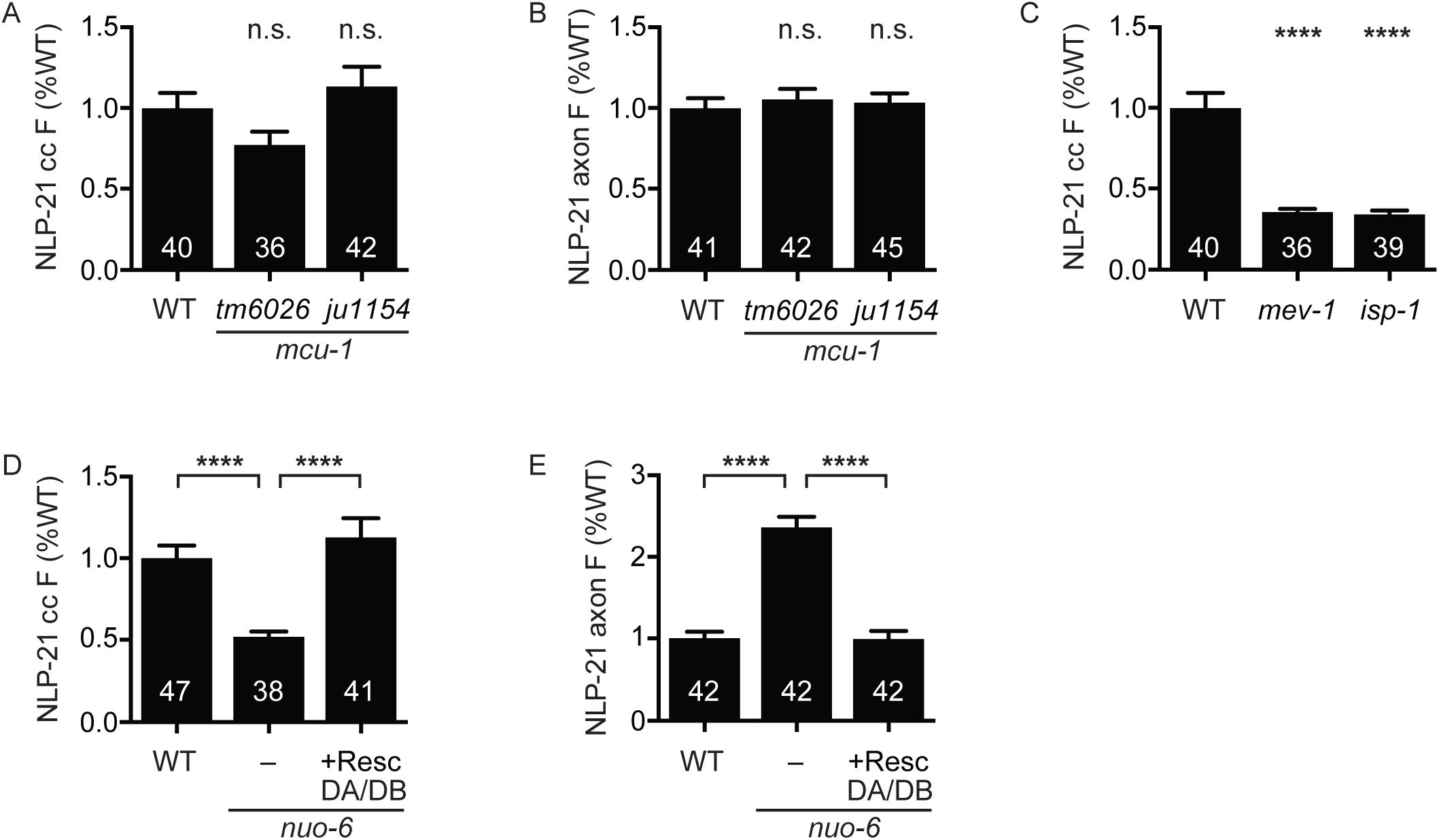
Neuropeptide release requires oxidative phosphorylation but not mitochondrial calcium uptake. (**A, B**) Mutation of the mitochondrial calcium uniporter *mcu-1* does not alter NLP-21 coelomocyte (**A**) or axonal (**B**) fluorescence. (**C-E**) Oxidative phosphorylation is required for neuropeptide secretion cell-autonomously. (**C**) Mutation of the complex II component *mev-1* or the complex III component *isp-1* reduces NLP-21 coelomocyte fluorescence. Mutation of the complex I component *nuo-6* decreases NLP-21 coelomocyte (**D**) and increases NLP-21 axonal fluorescence (**E**), both of which are rescued by expression of wildtype *nuo-6* transgene in cholinergic DA/DB motor neurons only. Number of animals analyzed is indicated for each genotype. Error bars indicate SEM. Values that differ significantly are indicated (****, p < 0.0001; n.s., not significant; Kruskal-Wallis test).

Next, we asked if neuropeptide secretion requires mitochondrial respiration. Mutations in the *nuo-6, isp-1*, and *mev-1* genes (which encode components of complex I, II, and III, respectively) reduce respiration rate (Ishii *et al*. 1998; Yang and Hekimi 2010b; Pfeiffer *et al*. 2011; Yee *et al*. 2014). These oxidative phosphorylation mutants also had decreased NLP-21 coelomocyte fluorescence, indicating decreased neuropeptide secretion (Fig. 2C, D). The *nuo-6* NLP-21 secretion defect was rescued by a transgene restoring NUO-6 expression specifically in the DA/DB neurons (Fig. 2D, E). These results support the idea that mitochondrial respiration acts in a cell-autonomous manner to promote neuropeptide secretion from DA/DB neurons.

### Increased mitochondrial ROS inhibits neuropeptide secretion

Disrupting oxidative phosphorylation can lead to reduced ATP levels and altered production of reactive oxygen species (ROS) (Senoo-Matsuda *et al*. 2001; Yang and Hekimi 2010a; Yee *et al*. 2014), both of which could affect neuropeptide secretion. Several steps in neurotransmission, such as establishment and maintenance of membrane potential and membrane fusion, are known to require ATP. Accordingly, we found that mutants of glycolytic enzymes also have decreased neuropeptide secretion (Fig. S1A).

Conflicting results have been reported for the effects of *mev-1*, *isp-1*, and *nuo-6* mutations on ATP levels (Senoo-MATSUDA *et al*. 2001; Yang and Hekimi 2010a; Yang and Hekimi 2010b; Yee *et al*. 2014). In addition, several findings suggest that neurotransmission may be more reliant on glycolysis than oxidative phosphorylation to meet its energy demands (Rangaraju *et al*. 2014; Jang *et al*. 2016; Ashrafi and Ryan 2017; Ashrafi *et al*. 2017). Prompted by these results, we investigated other potential mechanisms for neuropeptide secretion defects in the oxidative phosphorylation deficient mutants. Since *mev-1*, *isp-1*, and *nuo-6* mutants have all been shown to increase ROS levels (Ishii *et al*. 1998; SENOO-MATSUDA *et al*. 2001; Lee *et al*. 2010; Yang and Hekimi 2010a; Yang and Hekimi 2010b; Yee *et al*. 2014), we hypothesized that altered ROS production might contribute to decreased secretion in these mutants. To test this idea, we measured NLP-21 coelomocyte fluorescence in mutants lacking ROS detoxification enzymes. Superoxide dismutases (SODs) convert superoxide to hydrogen peroxide (H_2_O_2_), which can then be broken down to water by catalase (Fig. 3A). In *C. elegans*, the genes *sod-1* and *sod-5* encode the cytoplasmic Cu/ZnSODs (Larsen 1993; Jensen and Culotta 2005); *sod-2* and *sod-3* encode the mitochondrial MnSODs (Giglio *et al*. 1994; Suzuki *et al*. 1996; Hunter *et al*. 1997); and *sod-4* encodes an extracellular Cu/ZnSOD (Fujii *et al*. 1998). We analyzed NLP-21 secretion in all five SOD mutants (Fig. S1B, C). In *sod-*2 mutants (which lack the main mitochondrial SOD), NLP-21 secretion was significantly reduced, as indicated by decreased NLP-21 coelomocyte fluorescence and increased NLP-21 axonal punctal fluorescence, both of which were rescued by a transgene restoring *sod-2* expression in DA/DB neurons (Fig. 3B, C). Thus, *ric-7* and *sod-2* mutants exhibited very similar defects in NLP-21 secretion, suggesting that SOD-2 and RIC-7 function together to promote NLP-21 secretion. Consistent with this idea, we found that the NLP-21 coelomocyte fluorescence defect observed in *sod-2; ric-7* double mutants was not significantly different from that observed in *ric-7* single mutant (Fig. 3D). Collectively, these results suggest that perturbing mitochondrial ROS detoxification in axons inhibits NLP-21 secretion.

**Figure 3:**
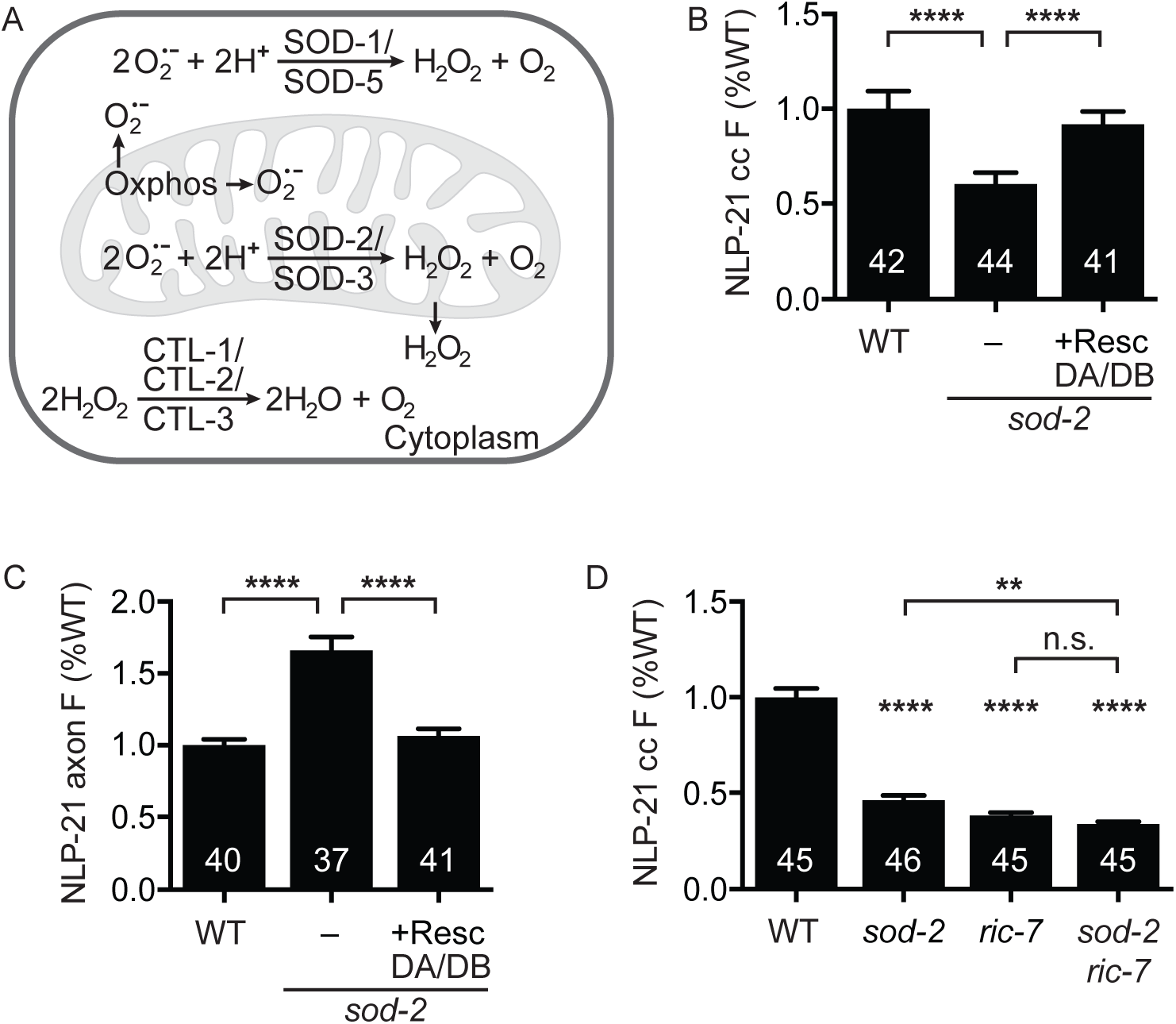
Mitochondrial superoxide detoxification is required for neuropeptide secretion. (**A**)Oxidative phosphorylation generates superoxide as by-products, which is converted to hydrogen peroxide by superoxide dismutases SOD-1 and SOD-5 in the cytoplasm and SOD-2 and SOD-3 in mitochondria. Hydrogen peroxide is membrane diffusible and is subsequently detoxified by catalases. (**B, C**) Mutation of *sod-2* decreases NLP-21 coelomocyte fluorescence (**B**) and increases NLP-21 axonal fluorescence (**C**), both of which are rescued by expressing a wildtype *sod-2* construct in cholinergic DA/DB motor neurons only. (**D**) NLP-21 coelomocyte fluorescence in wildtype, *sod-2* and *ric-7* single mutants, and *sod-2 ric-7* double mutant. In a *ric-7* mutant background, further mutation of *sod-2* has no effect. Number of animals analyzed is indicated for each genotype. Error bars indicate SEM. Values that differ significantly are indicated (****, p < 0.0001; **, p < 0.01; n.s., not significant; ANOVA (C) and Kruskal-Wallis test (B, D)).

To determine if increased cytoplasmic ROS levels regulate NLP-21 secretion, we analyzed *sod-1* mutants, which lack the major cytoplasmic SOD. In *sod-1* mutants, we expect increased cytoplasmic superoxide and decreased cytoplasmic H_2_O_2_. Unlike *sod-2* mutants, NLP-21 secretion was significantly increased in *sod-1* mutants (Fig. S1B). These results suggest that mitochondrial and cytoplasmic ROS have opposite effects on neuropeptide secretion. Taken together, these results support the idea that neuropeptide secretion is strongly inhibited by a variety of mitochondrial defects, including mutations disrupting axonal transport of mitochondria, mutations disrupting mitochondrial respiration, and mutations increasing mitochondrial ROS levels. Furthermore, in each case mitochondria act cell-autonomously to promote neuropeptide secretion.

### Hypoxia inhibits neuropeptide secretion

The neuropeptide secretion defects observed in mitochondrial mutants could result from decreased levels of ATP or other metabolites. Alternatively, decreased neuropeptide secretion could result from activation of stress response pathways. Several stress response pathways are induced in mitochondrial mutants, including those mediated by the hypoxia inducible factor (HIF-1), SKN-1/Nrf2, the mitochondrial unfolded protein response, and the AMP activated protein kinase (Sena and Chandel 2012; Hwang *et al*. 2014; Blackwell *et al*. 2015; Chang *et al*. 2017; Dues *et al*. 2017; Shpilka and Haynes 2018).

We focused on HIF-1 because *ric-7* mutants display hypoxic phenotypes (Jang *et al*. 2016) and mutations decreasing mitochondrial respiration stabilize HIF-1 (Lee *et al*. 2010; Hwang *et al*. 2014). Prompted by these results, we hypothesized that *ric-*7 mutants might undergo a hypoxic response that inhibits neuropeptide release. We did several experiments to test this idea. First, we asked if growth in reduced oxygen tension inhibits neuropeptide secretion. Consistent with this idea, growing wildtype animals in a hypoxic chamber (5 or 10% oxygen) for 24 hours significantly reduced NLP-21 coelomocyte fluorescence (Fig. 4A). By contrast, growth in hyperoxic conditions (100% oxygen) for the same duration had no effect (Fig. S2A). Thus, hypoxia was sufficient to inhibit NLP-21 secretion.

**Figure 4:**
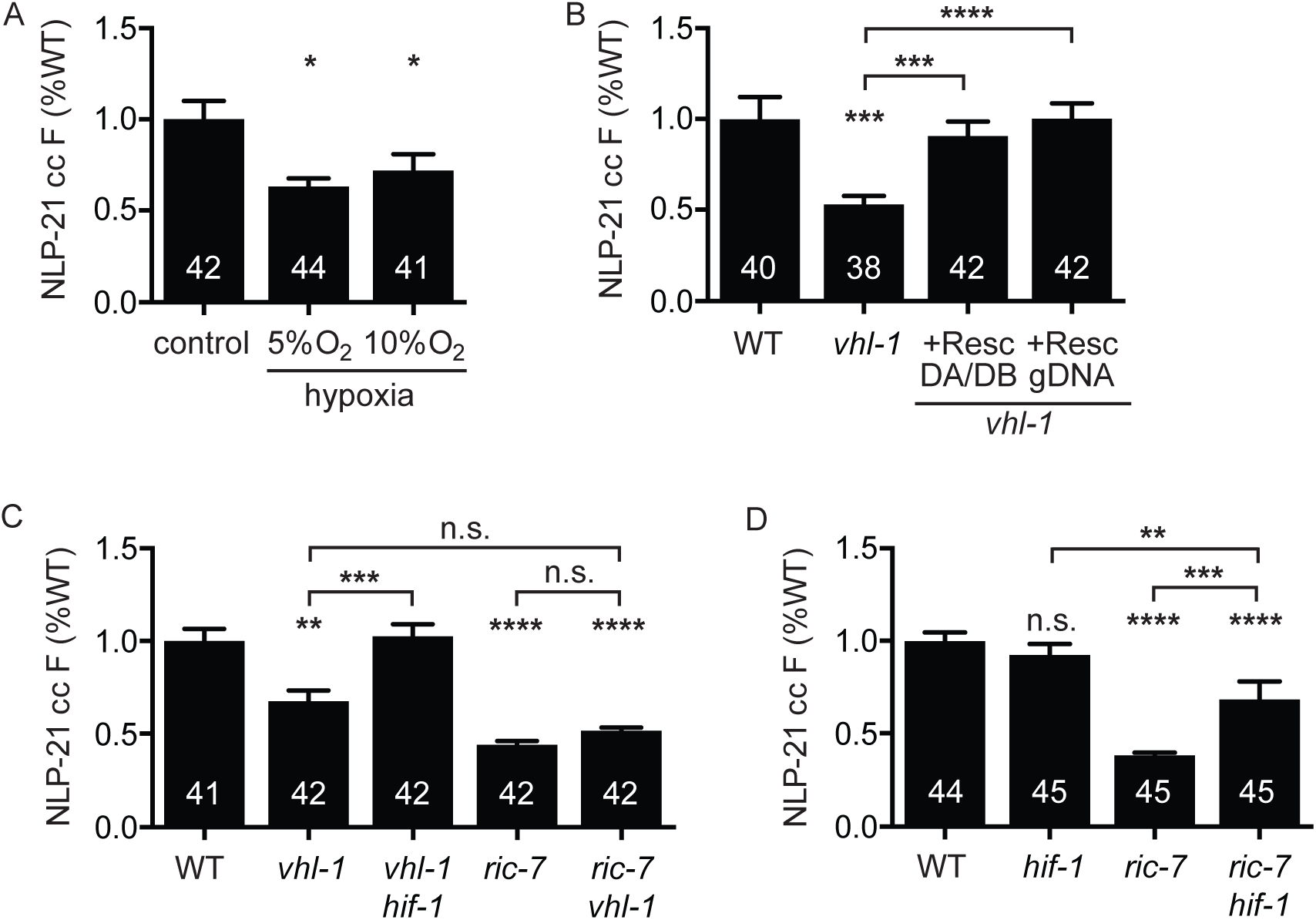
*ric-7* mutants inhibit neuropeptide secretion through activation of the hypoxic response. (**A**) NLP-21 coelomocyte fluorescence in worms that were either grown at atmospheric oxygen levels, at 5% oxygen, or at 10% oxygen for 24 hours. Worms grown under hypoxia show reduced NLP-21 coelomocyte fluorescence. (**B-D**) Comparison of NLP-21 coelomocyte fluorescence for the indicated genotypes. (**B**) NLP-21 coelomocyte fluorescence is reduced in *vhl-1* mutant worms and restored to normal levels by expressing a wildtype *vhl-1* rescue construct either under a DA/DB specific or the genomic *vhl-1* promoter. (**C**) Mutation of *hif-1* suppresses the decreased NLP-21 coelomocyte fluorescence of *vhl-1* mutation. *vhl-1* and *ric-7* mutations have no additive effect on NLP-21 coelomocyte fluorescence. (**D**) Mutation of *hif-1* partially restores NLP-21 coelomocyte fluorescence in *ric-7* mutants. Number of animals analyzed is indicated for each genotype. Error bars indicate SEM. Values that differ significantly are indicated (****, p < 0.0001; ***, p < 0.001; **, p < 0.01; *, p < 0.05; n.s., not significant; Kruskal-Wallis test).

Next, we asked if activation of HIF-1 was sufficient to inhibit NLP-21 secretion. To test this idea, we analyzed *vhl-1* mutants. VHL-1 is the *C. elegans* ortholog of the von Hippel Lindau tumor suppressor protein, which promotes HIF-1 ubiquitination under normoxic conditions. In *vhl-1* mutants, HIF-1 is stabilized and constitutively activates the hypoxic transcriptional response (Kaelin 2007). As expected, NLP-21 coelomocyte fluorescence was significantly decreased in *vhl-1* mutants and this defect was rescued by a transgene that restores VHL-1 expression specifically in the DA/DB neurons (Fig. 4B). The NLP-21 secretion defect in *vhl-1* mutants was abolished in *hif-1*; *vhl-1* double mutants, confirming that the *vhl-1* secretion defect was a consequence of stabilized HIF-1 (Fig. 4C). Taken together, these results suggest that activation of the HIF-1 hypoxia response in DA/DB neurons is sufficient to inhibit NLP-21 secretion.

Does activation of HIF-1 account for the *ric-7* neuropeptide secretion defect? We did two experiments to test this idea. First, if *ric-7* mutations inhibit neuropeptide secretion by activating the hypoxic response, then *hif-1* mutations should restore secretion in these mutants. As predicted, *ric-7 hif-1* double mutants had significantly higher NLP-21 coelomocyte fluorescence than *ric-7* single mutants (Fig. 4D). This effect was specific because inactivating two other stress-activated transcription factors (DAF-16/FOXO and SKN-1/Nrf) did not increase NLP-21 coelomocyte fluorescence in *ric-7* mutants (Fig. S2B, C). Second, if *ric-7* mutations inhibit neuropeptide secretion by stabilizing HIF-1, then *ric-7* and *vhl-1* mutations should not have additive effects on NLP-21 secretion in double mutants. Consistent with this idea, NLP-21 coelomocyte fluorescence in *vhl-1; ric-7* double mutants was not significantly different from that in either single mutant (Fig. 4C). By contrast, *vhl-1* and *unc-31*/CAPS mutations did have additive effects on NLP-21 secretion in double mutants (Fig. S2D). Collectively, these results support the idea that lack of axonal mitochondria (in *ric-7* mutants) elicits a cell-autonomous hypoxic response that stabilizes HIF-1 and inhibits neuropeptide secretion.

## DISCUSSION

Here, we show that axonal transport of mitochondria is critical for neuropeptide release and locomotion in *C. elegans*. Impaired mitochondrial transport, reduced oxidative phosphorylation rates, and increased mitochondrial ROS levels all inhibited neuropeptide secretion cell-autonomously, whereas interfering with mitochondrial calcium uptake had no effect. The effects of axonal mitochondria on neuropeptide release are mediated (at least in part) by changes in HIF-1 activity. Activating HIF-1 inhibited neuropeptide secretion while *hif-1* null mutations partially restored secretion in mutants lacking axonal mitochondria. These results demonstrate that modifications of mitochondrial transport or function lead to cell-autonomous changes in neuropeptide secretion, thus providing a mechanism whereby the metabolic or redox state of a neuron could alter the global behavioral state of the animal.

### Mitochondrial transport mutants as a model to identify mitochondrial function in neurotransmission

Blocking mitochondrial transport into axons can result in a variety of synaptic defects by a number of different mechanisms. For example, mouse syntabulin knockouts are defective for mitochondrial transport in cultured superior cervical ganglion neurons, which is accompanied by accelerated synaptic depression, slowed recovery after synaptic vesicle depletion, and impaired short-term plasticity (Ma *et al*. 2009). In contrast, decreased axonal mitochondria in rat syntaphilin mutant hippocampal neurons slow presynaptic calcium clearance and enhance short-term facilitation (Kang *et al*. 2008). Here, we find that the absence of axonal mitochondria induces a hypoxic response that inhibits neuropeptide secretion, thus identifying a new mechanism by which mitochondrial transport defects alter neurotransmission. Although *ric-7* is not conserved beyond nematodes and has no recognizable structural domains, it is conceivable that RIC-7 functions analogously to syntabulin (which is a KIF5B cargo adaptor for mitochondrial transport) (Ma *et al*. 2009). While it is likely that RIC-7 promotes transport of other unidentified cargoes, restoring axonal mitochondria in *ric-7* mutants (by expressing mtruck) rescued the NLP-21 secretion defects. Thus, mitochondrial transport defects alone account for the impaired neuropeptide secretion of *ric-7* mutants.

### Neuropeptide secretion does not require the mitochondrial calcium uniporter MCU-1

Mitochondrial calcium uptake regulates synaptic strength in a variety of systems (Werth and Thayer 1994; Tang and Zucker 1997; Billups and Forsythe 2002; Medler and Gleason 2002; David and Barrett 2003; Talbot *et al*. 2003; Kim *et al*. 2005; Chouhan *et al*. 2010; Shutov *et al*. 2013; Kwon *et al*. 2016). However, other studies reported limited or undetectable effects of mitochondrial calcium uptake on neurotransmitter release (Guo *et al*. 2005; Verstreken *et al*. 2005; Ma *et al*. 2009), indicating that mitochondrial effects on local calcium may very between cell types and systems. Using two independent *mcu-1* null alleles, we show that neuropeptide secretion from *C. elegans* motor neurons is independent of the mitochondrial uniporter. This suggests that other calcium clearance mechanisms (such as the endoplasmatic reticulum, Na^+^/Ca^2+^ pumps, or membrane Ca^2+^ ATPases) may have the dominating effect in DA/DB neurons (Rizzuto *et al*. 1998; Emptage *et al*. 2001; Liang *et al*. 2002; Bouchard *et al*. 2003; De JUAN-SANZ *et al*. 2017).

Three matrix dehydrogenases are activated by calcium ions, thus coupling mitochondrial calcium levels to bioenergetics and metabolic flux (Denton and Mccormack 1980). Accordingly, knockout of MCU attenuates metabolic rate in a variety of cell types (Kamer and Mootha 2015). One might therefore expect that *mcu-1* mutant animals should inhibit neuropeptide secretion indirectly due to changes in mitochondrial metabolism. However, previous studies reported that *mcu* knockouts in *C. elegans* are viable and fertile, while *mcu* knockout mice have reduced exercise capacity but have unaltered basal metabolic rates (Pan *et al*. 2013; Xu and Chisholm 2014). Mitochondrial calcium levels are strongly reduced (25% of wildtype levels) but not eliminated in *mcu* knockouts, implying that alternate mechanisms for calcium entry into mitochondria must exist (Feng *et al*. 2013; Murphy *et al*. 2014). The residual mitochondrial calcium levels may be sufficient to allow *mcu-1* mutant worms to sustain normal levels of neuropeptide secretion.

### Increased mitochondrial ROS inhibits neuropeptide secretion

Two findings in this study suggest that increased mitochondrial ROS inhibits neuropeptide secretion cell-autonomously. First, neuropeptide secretion is reduced in mutants lacking the major mitochondrial superoxide dismutase, *sod-2*. Second, mutations in the genes *nuo-6*, *mev-1*, and *isp-1* all lead to elevated mitochondrial ROS levels (Senoo-MATSUDA *et al*. 2001; Yang and Hekimi 2010a; Yang and Hekimi 2010b; Yee *et al*. 2014) and decreased neuropeptide secretion. Because of their chemical reactivity, ROS can damage lipids, proteins, and DNA (Cross *et al*. 1987), and oxidative stress has been associated with many neurodegenerative diseases. ROS can also function as a second messenger in signal transduction pathways by oxidizing critical thiols within proteins, thereby regulating numerous biological processes, including metabolic adaptation, differentiation, and proliferation (Schieber and Chandel 2014; Reczek and Chandel 2015). Intracellular ROS promotes activation of stress response pathways in distant tissues. For example, ROS-induced activation of the mitochondrial unfolded protein response (UPR^mito^) in multiple classes of *C. elegans* neurons elicits a stress response in the intestine (Berendzen *et al*. 2016; Shao *et al*. 2016). In this case, the non-autonomous effect of ROS was mediated by increased neuronal secretion of serotonin and FLP-2. By contrast, we observed decreased NLP-21 secretion in motor neurons when mitochondrial function was impaired. Taken together, these results suggest that mitochondria can either promote (FLP-2) or inhibit (NLP-21) neuropeptide secretion, depending on the activating stimulus and cell type.

One surprising observation was that mutations of *sod-1* and *sod-2* have opposite effects on neuropeptide secretion, although both lead to increased ROS levels (Van RAAMSDONK and Hekimi 2009; Yanase *et al*. 2009). Recent findings suggest that the location of ROS production can influence downstream events (Van RAAMSDONK 2015). For example, in a ROS-sensitized *clk-1* mutant background of *C. elegans*, further increase of cytoplasmic ROS by mutation of *sod-1* shortens lifespan whereas increase of mitochondrial ROS by *sod-2* mutation increases lifespan (Schaar *et al*. 2015). How different subcellular localizations of ROS produce distinct phenotypes is poorly understood. One possible scenario could be that accumulation of cytoplasmic ROS in *sod-1* mutants has a more direct and higher impact on other compartments, such as the endoplasmatic reticulum (ER), than the increase of mitochondrial ROS in *sod-2* mutants. Disrupting the ER’s redox state is likely to interfere with protein folding and, consequently, could activate the ER unfolded protein response (UPR^ER^), thereby altering expression of a broad group of genes. It will be interesting to determine if activating the UPR^ER^ also alters neuropeptide secretion in a cell-autonomous manner.

### Activation of the hypoxic response by multiple pathways

Acute neural activity relies on an increase in oxidative phosphorylation for ATP supply, which results in decreased oxygen levels (Hall *et al*. 2012; Rangaraju *et al*. 2014). Therefore, oxygen depletion and HIF-1 activation provide a way to sense and adapt to metabolic stress. At normal oxygen levels, HIF-1 is continually hydroxylated by oxygen-dependent prolyl hydroxlyases. Hydroxylated HIF-1 is ubiquitinated by an E3 ligase containing VHL-1, which results in HIF-1 degradation. During hypoxia, the prolyl hydroxylases are inactive and HIF-1 becomes stabilized, initiating transcription of a variety of genes to adapt to the hypoxic condition. Prior studies suggest that mutations altering mitochondrial respiration can also stabilize HIF-1 (Lee *et al*. 2010; Hwang *et al*. 2014). Mitochondria have been proposed to regulate HIF-1 stability through a variety of mechanisms. Mitochondrially produced ROS are proposed to directly activate HIF-1 (Chandel *et al*. 2000; Brunelle *et al*. 2005; Guzy *et al*. 2005; Mansfield *et al*. 2005). Other studies suggest that altered abundance of mitochondrial metabolites stabilize HIF-1 (Lu *et al*. 2005; Hewitson *et al*. 2007; Koivunen *et al*. 2007; Mishur *et al*. 2016). Therefore, HIF-1 could become activated in *ric-7* mutants because the absence of mitochondria results in elevated ROS production or altered abundance of mitochondrial metabolites. NLP-21 secretion was inhibited when HIF-1 was activated by *vhl-1* mutations, by hypoxic growth conditions, or by mutations altering mitochondrial respiration and ROS levels. Thus, our results are compatible with either mechanism.

Activating HIF-1 initiates a comprehensive transcriptional program that, among others, reduces oxidative phosphorylation and promotes glycolysis (Wang *et al*. 1995; Rankin and Giaccia 2016). Activated HIF-1 also promotes axon regeneration following injury (Nix *et al*. 2014; Cho *et al*. 2015; Alam *et al*. 2016), suggesting a neuroprotective role in certain conditions. This study identifies a new role for HIF-1 in inhibiting neuropeptide secretion, indicating that in addition to cell-autonomous metabolic rewiring, activation of HIF-1 might (directly or indirectly) alter intercellular signaling, thus facilitating the communication of cell-specific stresses to different cells and tissues to induce broad physiological changes.

## ACKNOWLEDGEMENTS

We thank the following for strains, advice, reagents, and comments on the manuscript: the *Caenorhabditis* Genetics Center (CGC), S. Mitani, A. Chisholm, and Y. Jin for strains, E. Jorgensen for the mtruck and tom-20 constructs, J. Meisel for assistance with hypoxia assays, and members of the Kaplan lab for comments on the manuscript. This work was supported by a Human Frontiers Post-doctoral fellowship to TZ (LT000234/2013), and by an NIH research grant to JMK (DK80215).

---

**Figure S1:**
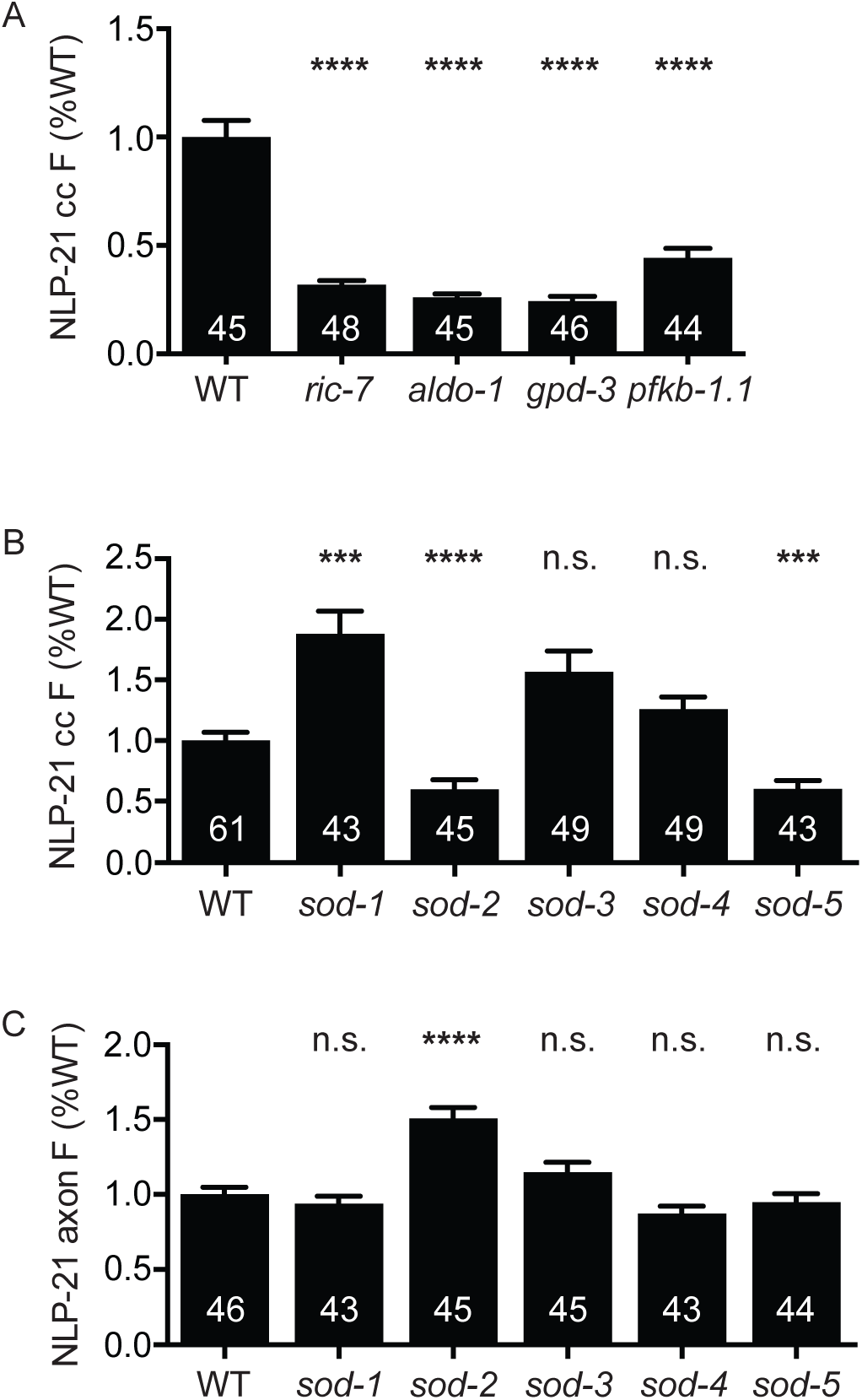
Neuropeptide secretion is sensitive to changes in ATP and reactive oxygen species. NLP-21 coelomocyte fluorescence (**A, B**) and axonal fluorescence (**C**) in mutants of glycolysis and superoxide dismutase genes. Number of animals analyzed is indicated for each genotype. Error bars indicate SEM. Values that differ significantly are indicated (****, p < 0.0001; ***, p < 0.001; n.s., not significant; Kruskal-Wallis test).

**Figure S2:**
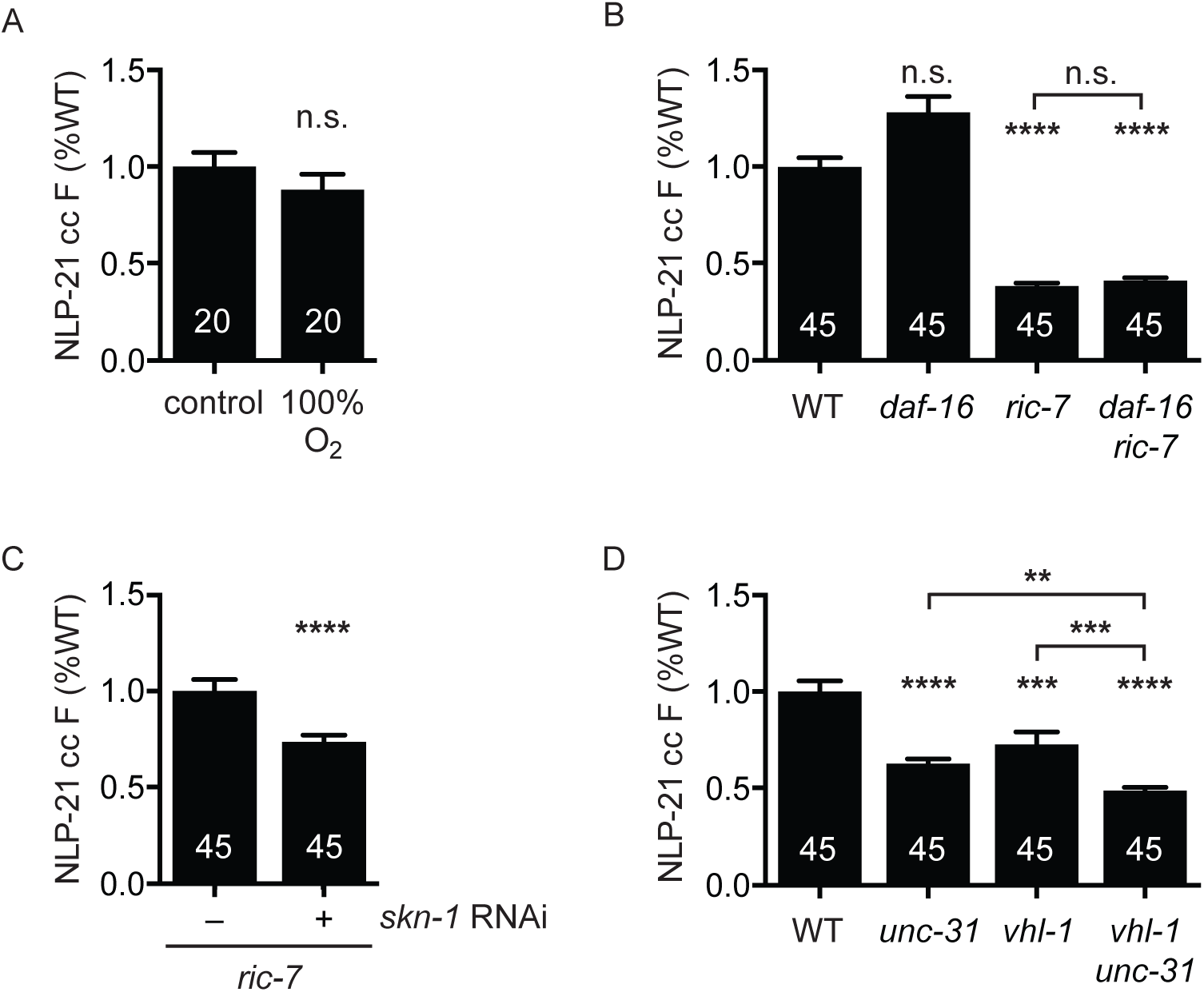
Neither hyperoxia nor activation of the DAF-16/FOXO or SKN-1/NRF stress responses can account for the neuropeptide secretion defect of *ric-7* mutants. (**A**) Comparison of NLP-21 coelomocyte fluorescence in worms grown at atmospheric oxygen levels and those grown at 100% oxygen for 24 hours. (**B**) Comparison of NLP-21 coelomocyte fluorescence for the indicated genotypes. Mutation of *daf-16* has no effect in wildtype or in a *ric-7* mutant background. (**C**) *skn-1* RNAi reduces NLP-21 coelomocyte fluorescence in *ric-7* mutants. (**D**) Comparison of NLP-21 coelomocyte fluorescence for the indicated genotypes. The NLP-21 coelomocyte fluorescence defects of *unc-31* and *vhl-1* mutations are additive. Number of animals analyzed is indicated for each genotype. Error bars indicate SEM. Values that differ significantly are indicated (****, p < 0.0001; ***, p < 0.001; n.s., not significant; Student’s t-test (A), Kruskal-Wallis test (B, D), and Mann-Whitney test (C)).

